# Sex differences in NK cells mediated by the X-linked epigenetic regulator UTX

**DOI:** 10.1101/2022.04.21.489076

**Authors:** Mandy I. Cheng, Luke Riggan, Joey H. Li, Rana Yakhshi Tafti, Scott Chin, Feiyang Ma, Matteo Pellegrini, Haley Hrncir, Arthur P. Arnold, Timothy E. O’Sullivan, Maureen A. Su

**Author notes:** Corresponding Authors: Timothy E. O’Sullivan, PhD, David Geffen School of Medicine at UCLA 615 Charles E. Young Drive South, BSRB 245F Los Angeles, CA 90095; Maureen A. Su, MD, David Geffen School of Medicine at UCLA 615 Charles E. Young Drive South, BSRB 290C Los Angeles, CA 90095.

## Abstract

Viral infection outcomes are sex-biased, with males generally more susceptible than females. Paradoxically, the numbers of anti-viral natural killer (NK) cells are increased in males compared to females. Using samples from mice and humans, we demonstrate that while numbers of male NK cells are increased compared to females, they display impaired production of the anti-viral cytokine IFN-γ. These sex differences were not due solely to divergent levels of gonadal hormones, since these differences persisted in gonadectomized mice. Instead, these differences can be attributed to lower male expression of X-linked *Kdm6a* (UTX), an epigenetic regulator which escapes X inactivation in female NK cells. NK cell-specific UTX deletion in females phenocopied multiple features of male NK cells, which include increased numbers and reduced IFN-γ production. Integrative ATAC-seq and RNA-seq analysis revealed a critical role for UTX in the regulation of chromatin accessibility and gene expression at loci important in NK cell homeostasis and effector function. Consequently, NK cell-intrinsic UTX levels are critical for optimal anti-viral immunity, since mice with NK cell-intrinsic UTX deficiency show increased lethality to mouse cytomegalovirus (MCMV) challenge. Taken together, these data implicate UTX as a critical molecular determinant of NK cell sex differences and suggest enhancing UTX function as a new strategy to boost endogenous NK cell anti-viral responses.

## Introduction

Evolutionarily conserved sex differences exist in both innate and adaptive immune responses^1, 2^. While males are less susceptible to autoimmunity, they also mount a limited anti-viral immune response compared to females^3^. For instance, males have a higher human cytomegalovirus (HCMV) burden after infection, suggesting increased susceptibility to viral threats^4^. This has also been recently illustrated with the COVID-19 pandemic, in which the strong male bias for severe disease has been postulated to reflect sex differences in immune responses^5^. Multiple studies in humans and mice have recently reported differences in immune cell distribution and/or function in males vs. females^6–10^. However, the molecular basis for these differences, and the mechanisms by which these differences influence disease outcomes, remain incompletely understood.

Sex differences in mammals are defined not only by divergent gonadal hormones, but also by sex chromosome dosage^1^. Expression of a subset of X-linked genes, for example, is higher in females (XX) than males (XY). While females undergo random X chromosome inactivation (XCI) to maintain similar levels of X-linked protein expression between sexes, XCI is incomplete, with 3-7% of X chromosome genes escaping inactivation in mice and 20-30% escaping inactivation in humans^11^. As such, differential levels of X-linked gene expression in females vs. males have been linked to sex differences in a wide range of conditions including neural tube defects^12^ and autoimmune disease^13^.

As circulating type 1 innate lymphocytes, NK cells serve as an early line of defense against herpesvirus family members^14^. The importance of NK cells in anti-viral immunity is illustrated in patients with defective NK cell numbers or functionality, who are highly susceptible to infection by herpesviruses, such as HCMV and Epstein-Barr virus (EBV)^15, 16^. Similarly, NK cells are required for the control of mouse cytomegalovirus (MCMV) and other viral infections^17–, 19^, as mice with either genetic deficiency in NK cell function or loss of NK cell numbers have a significant increase in viral titers and mortality following MCMV infection^20–25^. Thus, NK cells are critical in anti-viral immunity across species.

Given this role for NK cells, it was therefore unexpected that NK cells are increased in virus-susceptible males^6–10^. Beyond NK cell numbers, other previously unappreciated sexually dimorphic NK cell feature(s) may instead account for sex differences with viral infections. Here, we show that NK cells in males are simultaneously expanded in number and deficient in effector function across mice and humans. These sex differences in NK cell composition and function are not completely due to hormonal differences, since these differences persisted in gonadectomized mice. Through expression screening, we identified the epigenetic regulator UTX (encoded by gene *Kdm6a*) as an XCI escapee that is expressed at significantly lower levels in male NK cells across mice and humans. Conditional ablation of UTX in female mouse NK cells, which mimics lower UTX expression in male NK cells, recapitulated NK cell phenotypes associated with male sex, such as increased NK cell numbers and lower production of IFN-γ. Furthermore, parallel Assay for transposase-accessible chromatin using sequencing (ATAC-seq) and bulk RNA sequencing (RNA-seq) of WT and UTX^NKD^ NK cells revealed a critical role for UTX in regulating chromatin accessibility and transcription of gene clusters involved in NK cell fitness (*Bcl2*) and effector response (*Ifng and Csf2).* Notably, UTX^NKD^ mice had increased mortality in response to MCMV infection, suggesting a critical role for UTX in the production of optimal anti-viral effector responses. Ultimately, our findings demonstrate that divergent UTX levels underlie sex differences in NK cell homeostasis and effector function. Enhancing UTX function may therefore represent a novel strategy for optimizing NK cell-mediated anti-viral immunity.

## Results

### NK cells display sexually dimorphic phenotypes independent of gonadal sex hormones

Due to the critical role of NK cells in anti-viral immunity and increased male susceptibility to cytomegalovirus (CMV) and other herpesvirus family infections^4^, it was surprising that multiple studies have reported that human males display increased NK cell numbers^6–10^. A recent investigation examining spleens of C57BL/6 mice also reported increased numbers of NK cells in males vs. females^26^. Consistent with this, our data show that splenic NK cells are increased in frequency **(Fig. 1a,b)** and absolute numbers **(Fig. 1c)** in male C57BL/6 mice compared to females. These findings suggest that other sexually dimorphic features beyond NK cell numbers may account for increased male susceptibility to viral infections. In response to viral infection, NK cells are critical for early production of proinflammatory cytokines, particularly IFN-γ^27^. To test if sex differences exist in NK cell-intrinsic function, we compared effector cytokine production in NK cells from female vs. male mice *ex vivo*. Stimulation with IL-12 and IL-18, cytokines known to induce robust IFN-γ production by NK cells^28^, resulted in lower IFN-γ production by male NK cells **(Fig. 1d-f)**. Similarly, stimulation of activated human NK cells (TCRβ^-^ CD3^-^ CD56^+^) isolated from peripheral blood mononuclear cells (PBMCs) with IL-12 and K562 leukemia cells resulted in lower %IFN-γ^+^ **(Fig. 1g)** and IFN-γ MFI **(Fig. 1h)** in male compared to female NK cells. Thus, while NK cell numbers are increased, male NK cell effector function is consistently reduced in both mice and humans.

**Figure 1:**
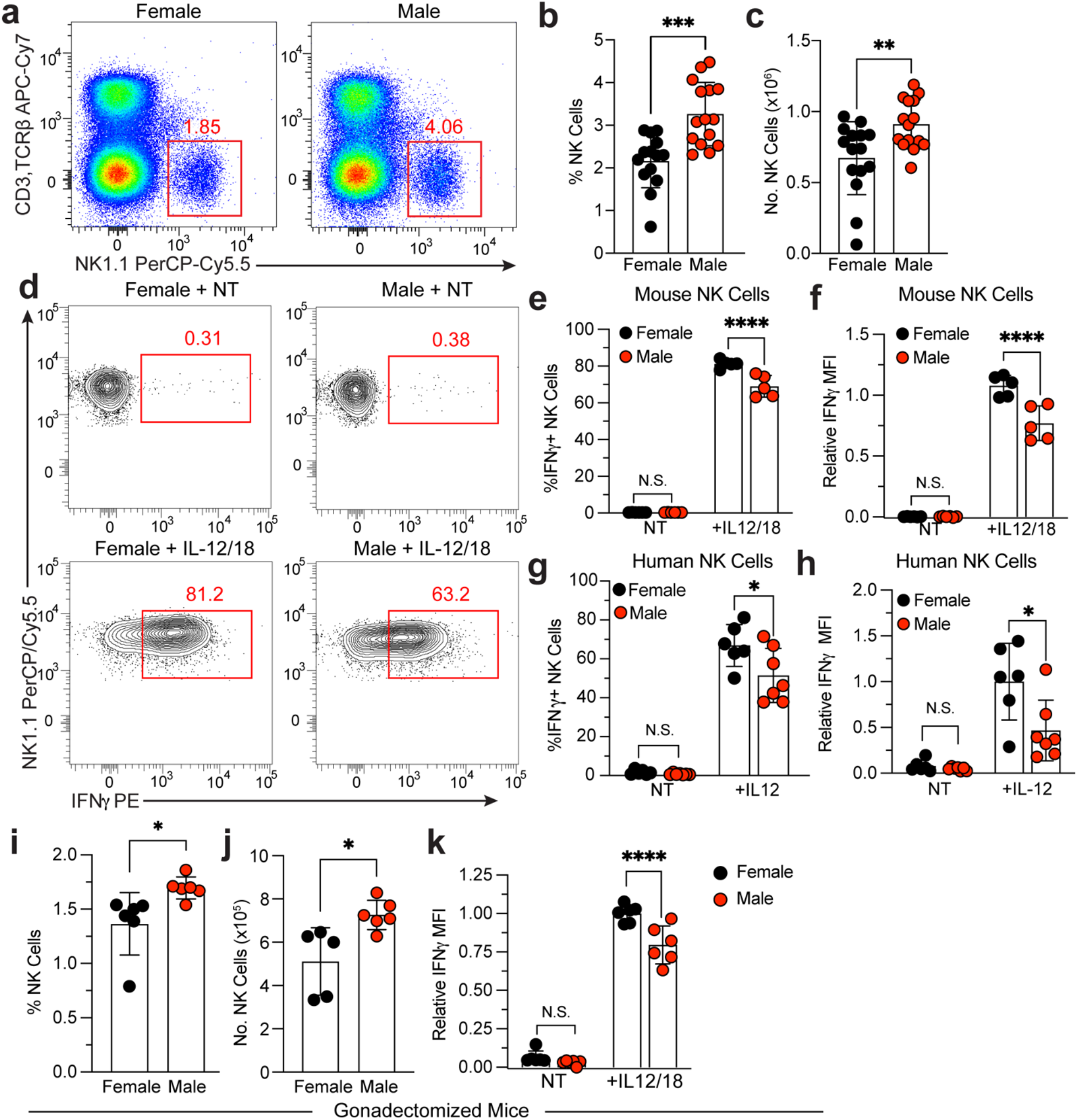
Sex differences in IFN-γ production and NK cell numbers in mouse and human NK cells are independent of gonadal hormones. **a)** Representative dot plots, **b)** percentage, and **c)** absolute numbers of splenic NK cells (CD3^-^ TCRβ^-^ NK1.1^+^) in female and male C57BL/6 mice (n = 15 per group). **d)** Representative contour plots, and bar graphs depicting **e)** percentages IFN-γ^+^ and **f)** relative IFN-γ mean fluorescence intensity (MFI) of total NK cells with no treatment (NT) or in response to IL-12 (20 ng/ml) and IL-18 (10 ng/ml) stimulation of splenic NK cells from female vs. male mice for 5 hours *ex vivo* (n = 5 per group, data representative of 2 independent experiments, normalized to female IL-12/18 treatment). **(g-h)** Human NK cells isolated from peripheral blood of female (n = 6) and male (n = 7) donors were cultured and stimulated with 10 ng/ml of IL-12 for 16 hours in the presence of K562 cells. **g)** Percentage of IFN-γ^+^ NK cells, and **h)** relative MFI of IFN-γ in TCRβ^-^CD3^-^CD56^+^ female and male human NK cells (normalized to female IL-12 treatment). **i)** Frequency and **j)** absolute numbers of splenic NK cells identified in gonadectomized mice (n = 6 per group). **k)** IFN-γ MFI of total splenic NK cells isolated gonadectomized female and male mice stimulated with IL-12 (20 ng/ml) and IL-18 (10 ng/ml) for 5 hours *ex vivo* (n = 6 per group). Samples were compared using two-tailed Student’s t test and data points are presented as individual mice with the mean ± SEM (N.S., Not Significant; *, p <0.05; **, p <0.01; ***, p<0.001; ****, p<0.0001).

Female or male sex is based on a composite of gonadal hormones (e.g., estrogens or androgens) and sex chromosomes (e.g., 46XX or 46XY)^1^. Previous studies demonstrated direct effects of gonadal hormones in regulation of IFN-γ production by NK cells^29^, but it remains possible that NK cell sex differences can also be attributed to sex chromosome complement. To test this possibility, we gonadectomized mice to abolish the effect of sex hormones. Gonadectomy failed to eliminate sex differences in NK cell frequency **(Fig. 1i),** absolute numbers **(Fig. 1j)** and IFN-γ protein production in response to cytokine stimulation **(Fig. 1k)**, indicating that gonadal hormones are not solely responsible for sex differences in NK cells. Thus, we hypothesized that chromosomal complement, in particular X chromosome dosage, may also play an important role.

### X-linked UTX escapes X-inactivation and has higher expression in female NK cells

While 46XX females undergo X chromosome inactivation (XCI) to control dosages of X-linked genes, a subset of genes escapes XCI (termed XCI escapees), often resulting in higher expression in females compared to males. Thus, XCI escapees are prime candidates for mediating phenotypic sex differences in NK cells. Five genes (*XIST*, *DDX3X, KDM6A, EIF2S3, KDM5C*) have previously been identified as XCI escapees in both humans and mice^30^. *XIST* was excluded from further analysis because it is not expressed in male cells due to its known role in X chromosome inactivation in female cells^1^. All 4 remaining genes were significantly downregulated in male vs. female NK cells, from humans **(Fig. 2a)** and mice **(Fig. 2b).** The greatest differential expression in both human and mouse NK cells was seen with *Kdm6a* (also known as UTX) **(Fig. 2a,b**). Male NK cells also expressed lower UTX protein levels compared to female NK cells in mice **(Fig. 2c,d)**. These data indicate that expression levels of *Kdm6a* (UTX) is sex-biased in NK cells.

**Figure 2:**
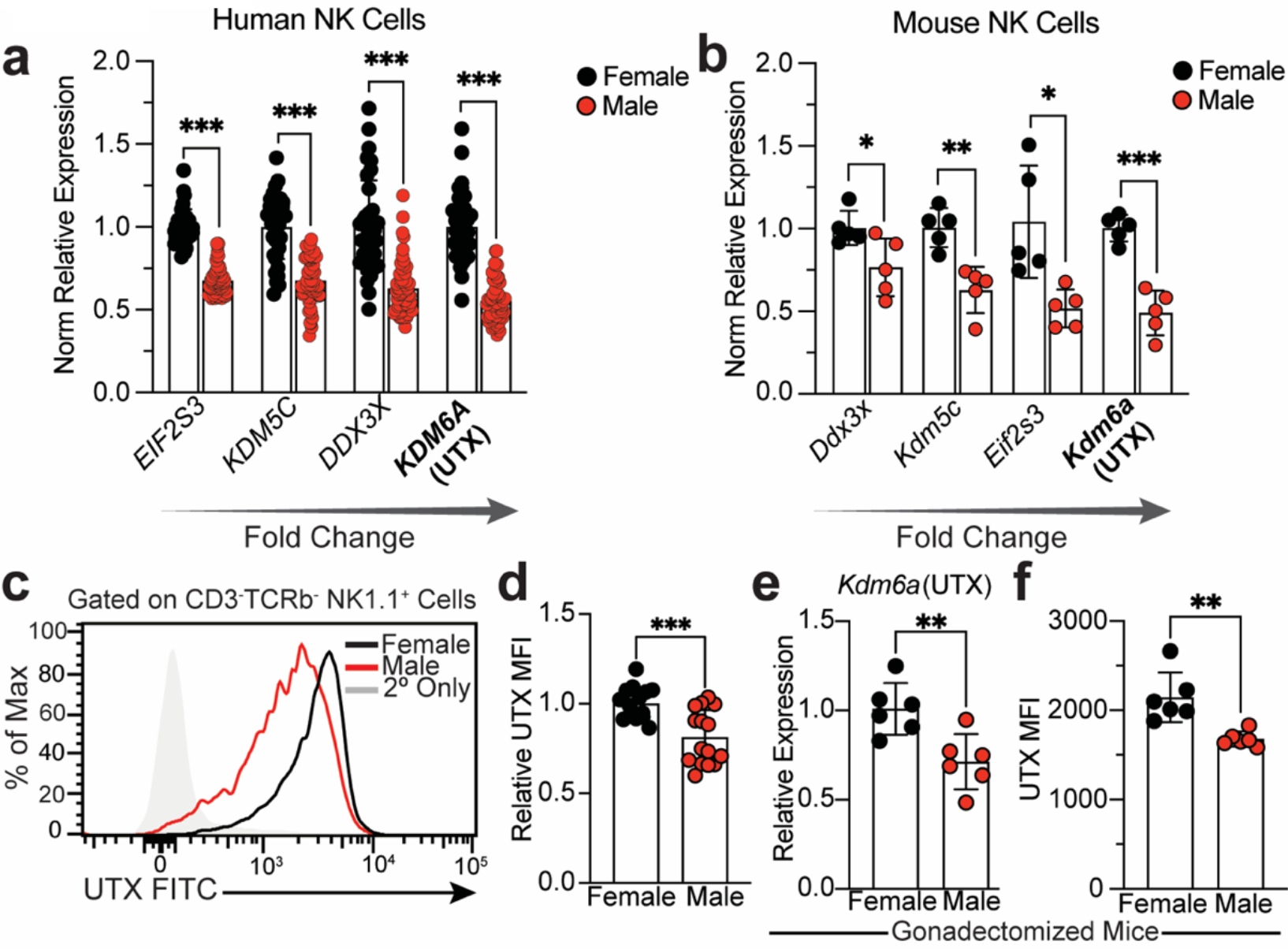
X-linked UTX displays sexually dimorphic gene expression independent of sex hormones. **a)** Relative expression of X-chromosome inactivation escapee genes using DICE database which performed RNA-seq on sorted NK cells from human females (n = 36) vs. males (n = 54) normalized to female. **b)** Relative expression of X chromosome inactivation escapee genes by RT-qPCR in NK cells from female vs. male mice (C57BL/6; 8 week old, n = 5 per group). Genes are ordered by increasing fold change. **c)** Representative histogram and **b)** relative MFI of UTX protein expression in splenic NK cells from naïve female vs. male mice by flow cytometry (C57BL/6; 8 week old). **e)** Relative expression of *Kdm6a* (UTX) by RT-qPCR of isolated splenic NK cells normalized to female and **f)** UTX MFI of NK cells by flow cytometry from spleens of gonadectomized female (ovariectomized) and male (castrated) mice (n = 6 per group). Samples were compared using two-tailed Student’s t test and data points are presented as individual mice with the mean ± SEM (*, p <0.05; **, p <0.01; ***, p<0.001).

In NK cells derived from gonadectomized mice, differences persisted in *Kdm6a* transcript levels **(Fig. 2e)** and UTX protein levels **(Fig. 2f).** Additionally, using the four core genotype (FCG) mouse model, which uncouples sex chromosome complement (XX or XY) and gonadal sex organ (ovaries or testes)^31^, *Kdm6a* transcript levels were also lower in mice with one X chromosome (XY) independent of gonadal composition **(Extended Data Fig. 2a)**. Together these findings suggest that increased UTX expression in female mice is not due to hormonal effects and instead point to a primary role for X chromosome dosage.

### UTX deletion recapitulates male NK cell phenotypes of frequency and IFN-γ production

To determine if UTX mediates sex differences in NK cells, we generated mice with a conditional deletion of UTX in NK cells (*Kdm6a*^fl/fl^ x *Ncr1*^Cre+^, referred to as UTX^NKD^ hereafter) with WT (*Kdm6a*^fl/fl^ x *Ncr1*^Cre-^) littermates used as controls. To control for gonadal hormone and sex chromosome effects, comparisons were made only in female mice (female WT vs. female UTX^NKD^ littermates). We confirmed decreased UTX protein expression in NK cells from UTX^NKD^ mice using flow cytometry **(Extended Data Fig. 2b,c).** Similar to male mice, female UTX^NKD^ mice displayed increased splenic NK cell frequency **(Fig. 3a,b)** and absolute numbers **(Fig. 3c)** in the spleen, blood, lungs, liver and bone marrow, demonstrating that this increase was not tissue specific. Furthermore, IFN-γ protein production in response to IL-12 and IL-18 stimulation was decreased in NK cells from UTX^NKD^ vs. WT mice **(Fig. 3d-f)**. These results implicate UTX in limiting NK cell numbers and promoting IFN-γ production, suggesting divergent UTX levels may play a causal role in NK cell sex differences.

**Figure 3:**
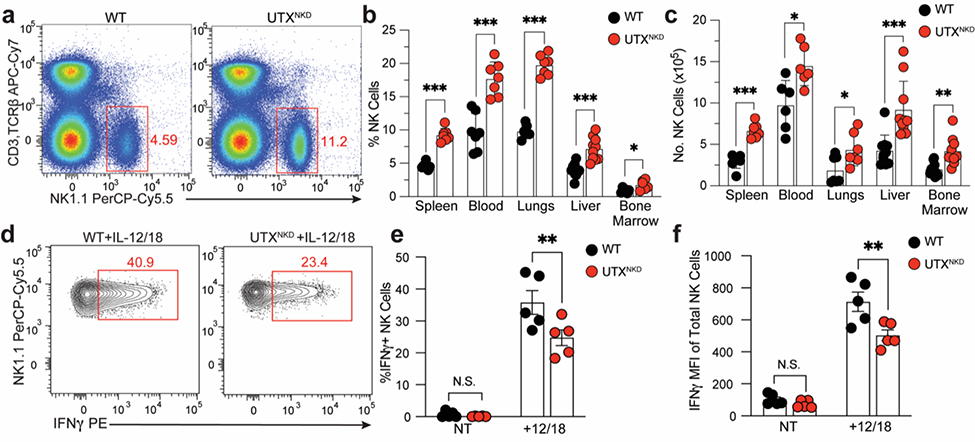
UTX regulates NK cell numbers and IFN-γ expression in NK cells. **a)** Representative flow cytometry dot plots of splenocytes from WT and UTX^NKD^ mice, gated on NK cells (CD3^-^,TCRβ^-^NK1.1^+^). **b)** Percentage and **c)** absolute numbers of NK cells isolated from the spleen, blood, lungs, liver, and bone marrow of WT and UTX^NKD^ mice (n = 6-10 per group). Numbers represent (spleen, lungs and liver – entire organ), (blood - per 1 ml of blood), and (Bone Marrow - 2 femurs per mouse). **d)** Representative contour plots, **e)** frequency, and **f)** MFI of IFN-γ of total NK cells in response to IL-12 (20 ng/ml) and IL-18 (10 ng/ml) stimulation of splenic NK cells from WT and UTX^NKD^ mice for 5 hours *ex vivo* (n = 5 per group). Data are representative of 2-4 independent experiments. Samples were compared using two-tailed Student’s t test and data points are presented as individual mice with the mean ± SEM (N.S., Not Significant; *, p <0.05; **, p <0.01; ***, p<0.001).

### Global changes in NK cell chromatin accessibility and transcription mediated by UTX

Recent studies have identified NK cell regulatory circuitry (regulomes) that prime innate lymphoid cells for swift effector responses even prior to NK cell activation^32, 33^. As an epigenetic modifier, UTX can alter transcription by organizing chromatin at regulatory elements of target gene loci^34^. To investigate the UTX-mediated modifications on chromatin accessibility and gene expression in NK cells, we performed Assay for Transposase-Accessible Chromatin using sequencing (ATAC-seq) in tandem with bulk RNA sequencing (RNA-seq) on sort-purified WT (CD45.1^+^) and UTX^NKD^ (CD45.2^+^) NK cells from WT:UTX^NKD^ mixed bone marrow chimeras (mBMCs) **(Fig. 4a)**. Using mBMCs allowed for an internally controlled experiment to minimize environmental confounding factors. Principle Component Analysis (PCA) of both ATAC-seq and RNA-seq data revealed that samples clustered together by genotype **(Fig. 4b)**, indicating that loss of UTX results in profound changes in both the chromatin landscape and transcriptome of NK cells. ATAC-seq revealed 3569 peaks decreased and 2113 peaks increased in accessibility in UTX^NKD^ compared to WT NK cells (Log_2_ Fold Change > ±0.5, adjusted p-value < 0.05, FDR < 0.05) **(Supplementary Table 1)**. Moreover, RNA-seq identified 577 decreased and 377 increased genes in UTX^NKD^ vs. WT (Log_2_ Fold Change > ± 0.5, adjusted p-value < 0.05, FDR < 0.05) **(Supplementary Table 2)**. Thus, these data suggest UTX plays an active role in controlling the NK regulome at baseline.

**Figure 4:**
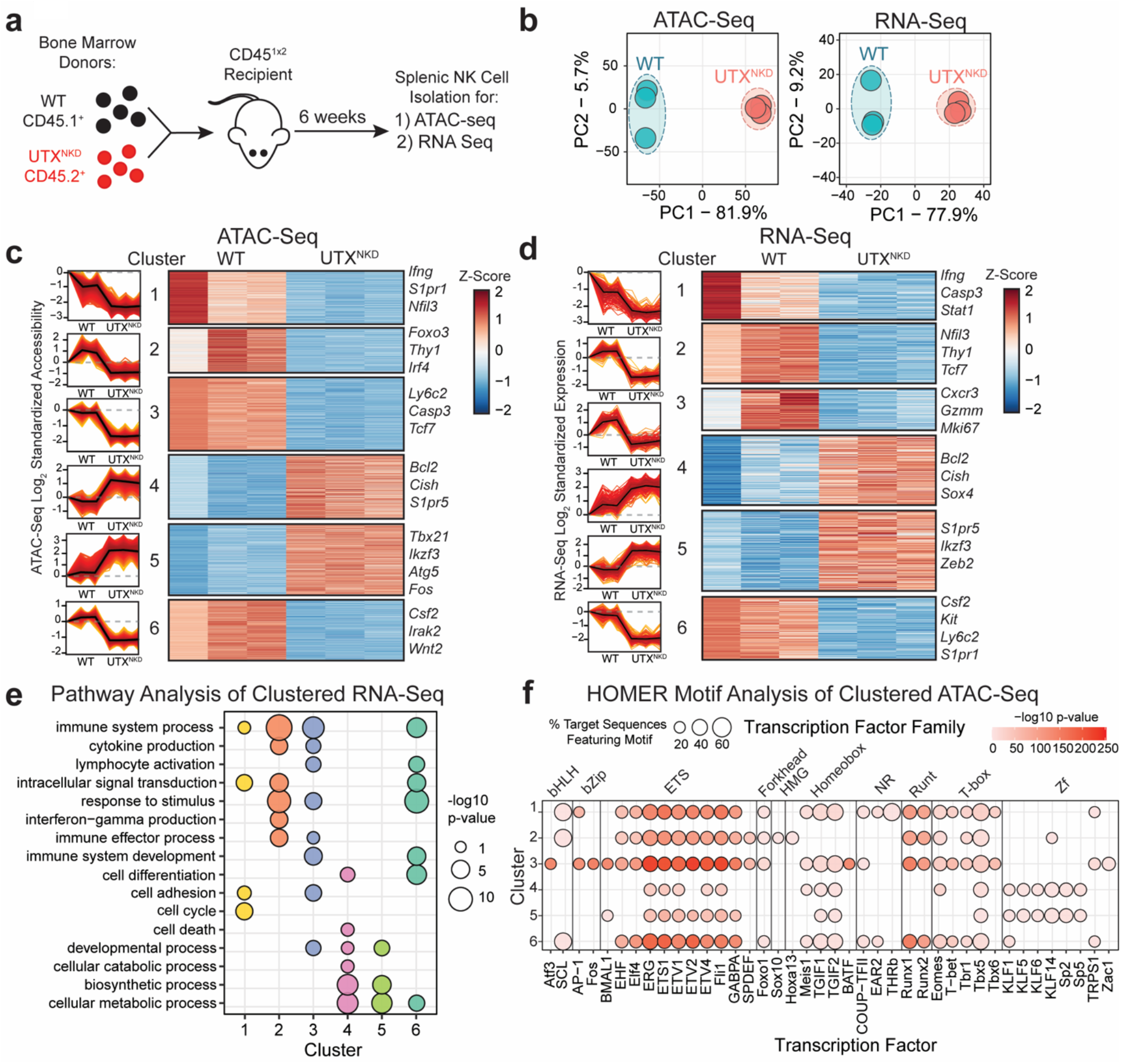
**Global changes in NK cell chromatin accessibility and transcription mediated by UTX. a**) Schematic of mixed bone marrow chimeras (mBMCs) produced by transferring cells from WT (CD45.1^+^) and UTX^NKD^ (CD45.2^+^) bone marrow donors into a lymphodepleted host (CD45^1×2^) and allowed to reconstitute for 6 weeks. Then NK cells were sorted from spleens of mBMCs for ATAC-seq and RNA-seq library preparation. **b)** Principal component analysis (PCA) of chromatin accessibility (ATAC-seq) and transcriptional (RNA-seq) changes in WT and UTX^NKD^ NK cells at steady state. **c,d)** Line graphs (left) and heatmap (right) of fuzzy c-means clustered differentially **c)** accessible peaks identified by ATAC-seq and **d)** expressed genes identified by RNA-seq of splenic NK cells from WT:UTX^NKD^ mBMCs (adjusted p-value < 0.05 and membership score > 0.5). Line graphs show mean (black line) and standard deviation (red ribbon) of mean-centered normalized log2 values of significant (FDR and adjusted p-value < 0.05) **c)** peaks of accessibility by ATAC-seq and **d)** gene expression by RNA-seq. **e)** Pathway analysis of significant fuzzy c-means clustered RNA-seq genes using g:Profiler with point size indicating −log10(p-value). **f)** HOMER motif analysis of significant fuzzy c-means clustered ATAC-seq peaks grouped by transcription factor family (top) and transcription factor (bottom). Point size indicates percentage of target sequences featuring motif and red gradient indicates - log10(p-value) of enrichment. All genes displayed in heatmaps met the following threshold of significance: FDR < 0.05, p < 0.05, and Log2FC > 0.5.

Integrative analysis of ATAC-seq and RNA-seq identified 400 genes that are both differentially accessible and expressed with a significant positive correlation (Spearman correlation: R = 0.62, p < 2.2×10^-16^) between the mean log_2_ fold change of ATAC-seq peaks and log_2_ fold change of RNA-seq expression **(Extended Data Fig. 3a)**. Fuzzy c-means clustering^35^ of both the ATAC-seq and RNA-seq datasets identified six major clusters which were significantly decreased (Clusters 1, 2, 3, and 6) or increased (Clusters 4 and 5) in accessibility (**Fig. 4c)** and expression **(Fig. 4d)** in UTX^NKD^ NK cells. For functional enrichment analysis, g:Profiler^36^ was used to analyze clusters of differentially expressed genes identified by RNA-seq **(Fig. 4e)**. Major pathways such as immune system process, cytokine production, IFN-γ production, lymphocyte activation, and immune effector process were associated with decreased expression in UTX^NKD^ (Clusters 1, 2, 3, and 6) **(Fig. 4e)**. At the same time, pathways such as developmental process, biosynthetic process, and metabolic process were significantly associated with increased expression in UTX^NKD^ (Clusters 4 and 5) **(Fig. 4e)**. Collectively, these findings implicate UTX in the coordinate regulation of genes associated with NK cell homeostasis and effector function.

Furthermore, UTX is known to interact with transcription factors (TFs) to coordinate target gene transcription^34^. To identify putative UTX TF partners, we performed HOMER (Hypergeometric Optimization of Motif Enrichment)^37^ TF motif analysis on each cluster of significant differentially accessible peaks **(Fig. 4f)**. TFs associated with modulating effector function during viral infection such as Runt (Runx1 and Runx2)^38^ and T-box (Eomes, T-bet, Tbr1 and Tbx6)^39^ family TFs were more significant and had a higher percentage of target motifs associated with clusters displaying decreased accessibility in UTX^NKD^ (Clusters 1, 2, 3, and 6) **(Fig. 4f)**. Conversely, TFs associated with proliferation, differentiation, and metabolism in the zinc finger family TFs (KLF1, KLF5, KLF6, KLF14, Sp2 and Sp5)^40^ were more significantly associated with clusters displaying increased accessibility (Clusters 4 and 5) **(Fig. 4f)**. These data suggest that UTX poises the chromatin accessibility of several genes at steady state known to influence NK cell fitness and effector responses, while also controlling genome-wide accessibility of transcription factor binding sites implicated in these processes.

### UTX coordinately regulates chromatin accessibility and expression of apoptosis pathway genes

The observed expansion of NK cell numbers in UTX^NKD^ mice **(Fig. 3a-c)** could either be due to higher NK cell proliferation or increased resistance to apoptosis. However, UTX^NKD^ NK cells paradoxically displayed a lower proportion of cells expressing the cell division marker Ki67 **(Extended Data Fig. 4a).** UTX^NKD^ NK cells also showed less CFSE dilution in response to IL-15, a cytokine known to induce NK cell proliferation **(Extended Data Fig. 4b,c)**. Thus, these results suggest that higher NK cell numbers observed in UTX^NKD^ mice were not due to increased proliferation. In contrast, increased survival is likely the cause of expanded NK cell numbers in UTX^NKD^ mice.

Among the differentially accessible and expressed genes in NK cells lacking UTX were those important in controlling apoptosis. An anti-apoptotic gene, *Bcl2,* showed increased accessibility and expression in UTX^NKD^ vs. WT NK cells (**Fig.5a, Extended Data Fig. 3b**).

**Figure 5:**
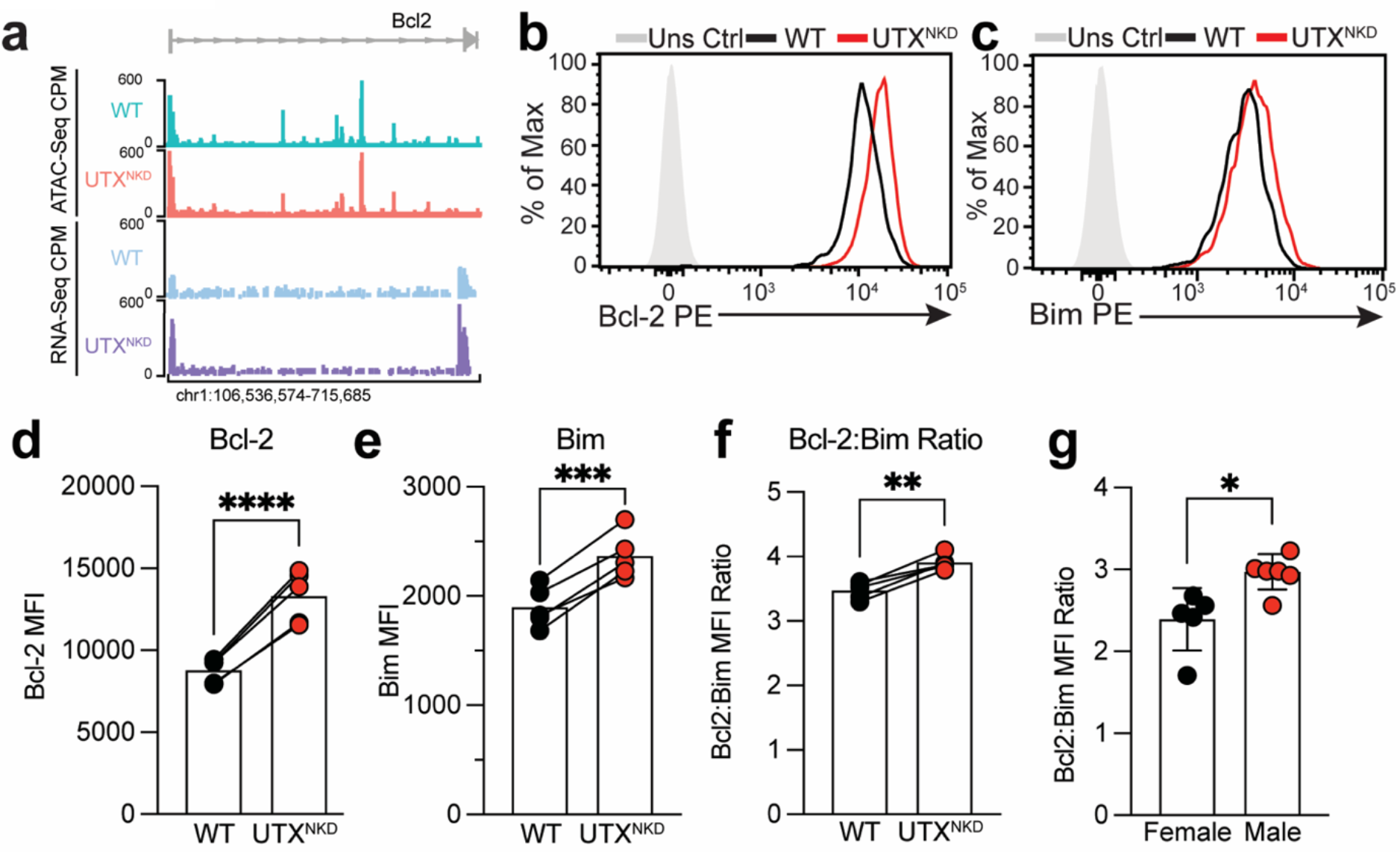
UTX regulates genes involved in apoptosis in NK cells through chromatin remodeling. **a)** Representative tracks of ATAC-seq signal (WT: green and UTX^NKD^: red) and RNA-seq transcript signal (WT: blue and UTX^NKD^: purple) of *Bcl2* in NK cells isolated at rest from WT:UTX^NKD^ mBMCs. Representative histogram of **b)** Bcl-2 and **c)** Bim protein expression, MFI of **d)** Bcl-2, **e)** Bim, and **f)** Bcl-2:Bim ratio from flow cytometric analysis of total peripheral blood NK cells (CD3^-^TCRβ^-^NK1.1^+^) from WT:UTX^NKD^ mBMCs (n = 4). **g)** Bcl-2:Bim MFI ratio of total splenic NK cells (CD3^-^TCRβ^-^NK1.1^+^) from gonadectomized female and male mice (n = 6 per group). Data represent 2-3 independent experiments. Samples were compared using two-tailed Student’s t test and data points are presented as individual mice with the mean ± SEM (*, p <0.05; **, p <0.01; ***, p<0.001; ****, p<0.0001).

Previous studies in mice have demonstrated that Bcl-2 can inhibit the pro-apoptotic function of Bim to promote NK cell survival^41^, thus, we interrogated the expression of these proteins in UTX^NKD^ NK cells. While naïve UTX^NKD^ NK cells showed increased intracellular protein expression of Bcl-2 compared to WT NK cells **(Fig. 5b,d),** UTX-deficient NK cells also displayed a modest increase in intracellular Bim levels **(Fig. 5c,e)**. Importantly, the Bcl-2:Bim ratio was significantly higher in UTX^NKD^ NK cells **(Fig. 5f)**, suggesting UTX-deficiency likely results in a higher proportion of pro-survival proteins present in NK cells. Notably, male NK cells also displayed a significant increase in Bcl-2:Bim ratio, which may underlie the expanded NK cell numbers observed in male mice **(Fig. 5g)**. These results implicate UTX in regulation of NK cell fitness to restrict numbers at homeostasis.

### UTX is critical for NK cell IFN-γ production and effector function

Since NK cells are early responders to MCMV, rapid effector molecule production is critical for NK cell mediated anti-viral control. ATAC-seq and RNA-seq of NK cells from WT:UTX^NKD^ mBMCs identified various chromatin accessibility and transcript changes at steady state **(Fig. 4b,c)**. Specifically, genes involved in the NK effector response (*Ifng* and *Csf2)*^27, 42^ showed decreased accessibility and expression in UTX^NKD^ compared to WT NK cells **(Fig. 6a, Extended Data Fig. 3b-d)**. Moreover, qRT-PCR verified decreases in *Ifng* transcript levels in UTX^NKD^ vs. WT NK cells at rest **(Fig. 6b).** Similarly, male NK cells, which express lower levels of UTX **(Fig.2c,d),** also had a similar decrease in *Ifng* transcript levels at rest **(Fig. 6c)**. Thus, through shaping of the chromatin landscape, UTX controls levels of effector gene transcripts available prior to immune challenge.

**Figure 6:**
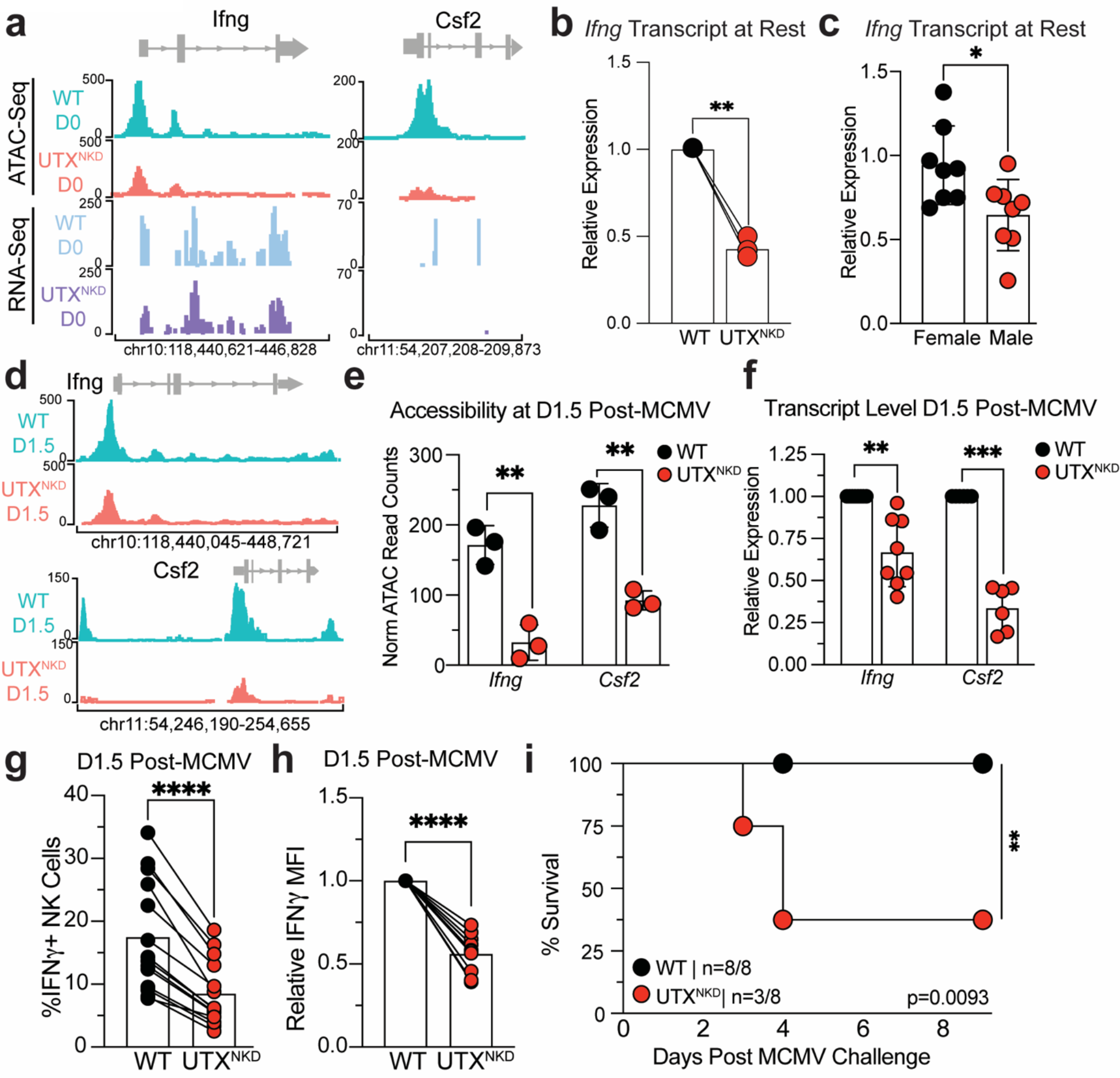
UTX controls the expression of effector genes critical for NK cell-mediated anti-viral immunity. **a)** Representative tracks of ATAC-seq signal (WT: green and UTX^NKD^: red) and RNA-seq transcript signal (WT: blue and UTX^NKD^: purple) of genes involved in NK effector function (*Ifng* and *Csf2)* in NK cells isolated at rest (D0) from WT:UTX^NKD^ mBMCs. **b)** Relative expression of *Ifng* by RT-qPCR of isolated splenic NK cells normalized to WT from WT:UTX^NKD^ mBMCs (n = 3). **c)** Relative expression of *Ifng* transcripts by RT-qPCR in NK cells from female vs. male mice normalized to female mice (C57BL/6; 8 week old, n = 8 per group). **d)** Representative tracks of ATAC-seq signal (WT: green and UTX^NKD^: red) of genes involved in NK effector function (*Ifng* and *Csf2)* in NK cells isolated at from WT:UTX^NKD^ mBMCs at D1.5 post-MCMV infection. **e)** Normalized read counts of accessibility at a representative peak located in the *Ifng* and *Csf2* loci at D1.5 post-MCMV infection of WT:UTX^NKD^ mBMCs (n = 3). **f)** Expression by RT-qPCR of *Ifng* and *Csf2* transcript levels at D1.5 post-MCMV infection of WT:UTX^NKD^ mBMCs relative to WT (n = 6-8). Flow cytometric analysis of **g)** percent IFN-γ^+^ and **h)** normalized IFN-γ MFI relative to WT in splenic NK cells at D1.5 Post MCMV infection of WT:UTX^NKD^ mBMCs (n = 14).**i)** Kaplan-Meier survival curves of WT and UTX^NKD^ mice infected with MCMV (n = 8 per genotype). Mantel-Cox test (**, p=0.0093). Data are representative of 2-3 independent experiments. Two-tailed paired Student’s t-test was performed, and data points depict individual mice with the mean ± SEM (*, p <0.05; **, p <0.01; ***, p<0.001; ****, p<0.0001).

To determine whether UTX deficiency also led to decreased chromatin accessibility and transcription of effector genes in NK cells during infection, we performed ATAC-seq on NK cells isolated from MCMV-infected WT:UTX^NKD^ mBMCs on D1.5 post-infection (PI). Similar to naïve UTX-deficient NK cells, D1.5 PI UTX^NKD^ NK cells also showed decreased chromatin accessibility at the *Ifng* and *Csf2* loci **(Fig. 6d,e, Extended Data Fig. 3e,f)**. qRT-PCR confirmed decreased *Ifng* and *Csf2* transcripts in NK cells from UTX^NKD^ mice at D1.5 PI **(Fig. 6f)**. Similarly, UTX^NKD^ NK cells showed decreased IFN-γ protein expression in UTX^NKD^ NK cells on D1.5 PI indicating that UTX expression in NK cells is required for optimal IFN-γ production following viral infection **(Fig. 6g,h)**. To confirm whether dosage of UTX expression in mature NK cells associates with NK cell production of IFN-γ during viral infection, we generated transgenic mice to achieve tamoxifen-inducible UTX deletion (*Rosa26*^ERT2CRE/+^ x *Kdm6a*^fl/fl^: iUTX^-/-^). Tamoxifen administration in WT:iUTX^-/-^ mBMCs resulted in differential degrees of UTX protein loss, and displayed a positive correlation between intracellular UTX levels and IFN-γ production on D1.5 PI **(Extended Data Fig. 5).** Since IFN-γ production by NK cells is critical for protection against MCMV^27^, we challenged WT and UTX^NKD^ mice with a sublethal dose of MCMV and monitored survival. While WT mice controlled MCMV infection (n = 8/8 survived), UTX^NKD^ mice rapidly succumbed to infection (n = 3/8 survived) **(Fig. 6i)**. These results demonstrate a requirement for NK cell-intrinsic UTX in the control of effector molecule production and protection against MCMV infection.

## Discussion

Sex is a critical biological variable in determining outcomes to viral infections^3^. This was recently illustrated with COVID-19, in which male sex was identified as a major risk factor for severe disease^5^. Moreover, recent studies have linked NK cell dysfunction within severe COVID-19 disease^43^. Given the importance of NK cells in anti-viral immunity, understanding the root causes of sex differences in NK cell biology will have far-reaching implications in optimizing anti-viral immunity. In this study, we demonstrated that lower expression of UTX in XX UTX^NKD^ NK cells mimics levels in XY NK cells, which may contribute to increased NK cell numbers and decreased IFN-γ production in males **(Extended Data Fig. 6a)**. UTX is expressed at lower levels in male NK cells across mice and human and this observation is independent of gonadal hormones in mice. NK cell UTX is required for controlling NK cell fitness, modulating accessibility of transcription factor binding motifs, increasing chromatin accessibility at effector gene loci, and poising NK cells for rapid response to virus infection **(Extended Data Fig. 6b)**. Together, these findings support a model in which divergent UTX expression contributes to sex differences in NK cell numbers and effector function.

Our findings indicate that UTX restricts NK cell numbers at steady state, since NK cells are increased at baseline in UTX^NKD^ mice. This is in contrast to UTX deficiency in other immune cell types, which have been reported to result in moderate (CD8^+^ and CD4^+^ T cells) or severe (iNKT) decreases in peripheral cell numbers^44–46^. Interestingly, T cell-specific UTX-deficiency is associated with CD8^+^ T cell accumulation during viral infection. Thus, it is possible that UTX-mediated gene programs that inhibit CD8^+^ T cell numbers during inflammation are shared by NK cells at rest^46^. Indeed, increased Bcl-2 levels were observed in both UTX-deficient NK cells and UTX-deficient CD8^+^ T cells, suggesting that UTX down-regulates this anti-apoptotic factor in both innate and adaptive cytotoxic lymphocytes.

NK cell-mediated effector functions during viral infection include cytokine production (IFN-γ) and cytotoxic molecule expression^47^. Our results from simultaneous ATAC-seq and RNA-seq suggest that UTX poises the chromatin landscape of NK cells at rest to quickly respond to viral challenge by increasing accessibility and transcription of effector loci such as *Ifng* and *Csf2* prior to viral infection^27, 42^. Decreased IFN-γ protein levels were seen at D1.5 post MCMV-infection in UTX-deficient NK cells, suggesting decreased effector functionality during inflammation. These results support a previous study that suggests the existence of an NK cell regulome^27, 42^, in which NK cell chromatin accessibility is actively maintained at steady-state and demonstrates a critical role for UTX in its maintenance before and during inflammation.

As a histone demethylase, UTX has intrinsic catalytic ability to demethylate H3K27me3 (a repressive histone mark) to poise chromatin for active gene expression^48^. In addition to its catalytic activity, UTX functions in multiprotein complexes with other epigenetic regulators (e.g. SWI/SNF, MLL4/5 and p300) to mediate chromatin remodeling in a demethylase-independent manner^48, 49^. In CD8^+^ T cells, UTX binds to enhancer and TSS of effector genes to promote effector gene programs in a demethylase-independent manner^46^. In contrast, demethylase activity in iNKT cells is required for the development and function and H3K27 methylation correlated with gene programs important for CD4^+^ T follicular helper cell development^44, 50^. Thus, the molecular mechanisms by which UTX functions appears to be immune cell specific. A previous study treated human NK cells with a small molecule inhibitor of H3K27me3 demethylases (GSK-J4) and found reduced cytokine expression (IFN-γ, TNF-α, GM-CSF, and IL-10) in response to *in vitro* stimulation^51^. However, GSK-J4 is not a specific tool for studying UTX-mediated mechanisms because it also inhibits Jmjd3, another H3K27me3 demethylase as well as having non-specific effects on other histone demethylases^52^. Thus, further studies using more precise genetic modification strategies are needed to understand the mechanisms by which UTX functions in NK cells.

UTX has been reported to interact with lineage specific TFs in T cells to target particular effector loci^34^. Our studies using HOMER motif analysis revealed potential interactions between UTX and TFs associated with modulating NK cell effector function during viral infection (Runx1, Runx2, Eomes, T-bet, Tbr1, and Tbx6). Moreover, these analyses also point to UTX interactions with TFs associated with NK cell proliferation, differentiation, and metabolism (KLF1, KLF5, KLF6, KLF14, Sp2 and Sp5). In line with a physiologic role for TFs in UTX-mediated control of chromatin accessibility, TF binding motifs were strongly enriched in ATAC-seq gene clusters. Further studies will be needed to experimentally verify these interactions in mouse and human NK cells.

NK cells are critical for control of HCMV infection in humans because NK cell-deficient individuals develop disseminated herpesvirus infections^16, 53^. Sex differences in immune response to multiple viruses have been reported, including immune responses to HCMV^53^. Our data support a model in which sex differences in anti-viral immunity can partially be explained by differences in UTX expression in NK cells, given the increased susceptibility of UTX^NKD^ mice to MCMV challenge. UTX deficiency has also been associated with Kabuki Syndrome and Turner Syndrome^44, 54^, two human conditions associated with immune dysregulation and increased infections. Our findings suggest the possibility that UTX deficiency in human NK cells may contribute to decreased viral immunosurveillance observed in these patients, although future work will be needed to support this hypothesis.

Weighing factors that define patient subsets with different immune responses will allow us to move past a “one-size-fits-all” therapeutic approach to a precision medicine paradigm. Understanding sex differences in NK cell function and their molecular underpinnings is an important step toward incorporating sex as a biological factor in treatment decisions. In males with severe viral illness, for instance, enhancing NK cell UTX activity may provide therapeutic benefit. We expect that these insights will be important not only in the setting of viral infections, but also in other infections and cancer, where NK cells also play an important role. These findings may also have important implications for adoptive cellular therapies, in which NK cells are the subject of intense interest^55, 56^.

## Acknowledgements

We thank members of the O’Sullivan and Su labs for helpful discussion. We thank the UCLA Technology Center for Genomics and Bioinformatics for RNA sequencing library preparation and the Cedars Sinai Applied Genomics, Computation, and Translational Core Facility for ATAC sequencing library preparation. T.E.O. is supported by the NIH (AI145997) and UC CRCC (CRN-20-637105). M.A.S. is supported by the NIH (NS107851, AI143894, DK119445) Department of Defense (USAMRAA PR200530), and National Organization of Rare Diseases. M.I.C. is supported by Ruth L. Kirschstein National Research Service Awards (GM007185 and AI007323), and Whitcome Fellowship from the Molecular Biology Institute at UCLA. L.R. is supported by the Warsaw fellowship from the MIMG department at UCLA. J.H.L. is supported by the NIH NIAMS (T32AR071307). A.P.A. is supported by NIH HD100298.

## Author Contributions

M.I.C., L.R., J.H.L., R.Y.T., H.H., A.P.A., T.E.O. and M.A.S. designed the study; M.I.C., J.H.L., R.Y.T., L.R., and S.C. performed the experiments; M.I.C., R.T.Y., F.M., and M.P. performed bioinformatics analysis; M.A.S, M.I.C. and T.E.O. wrote the manuscript.

---

**Extended Data Figure 1:**
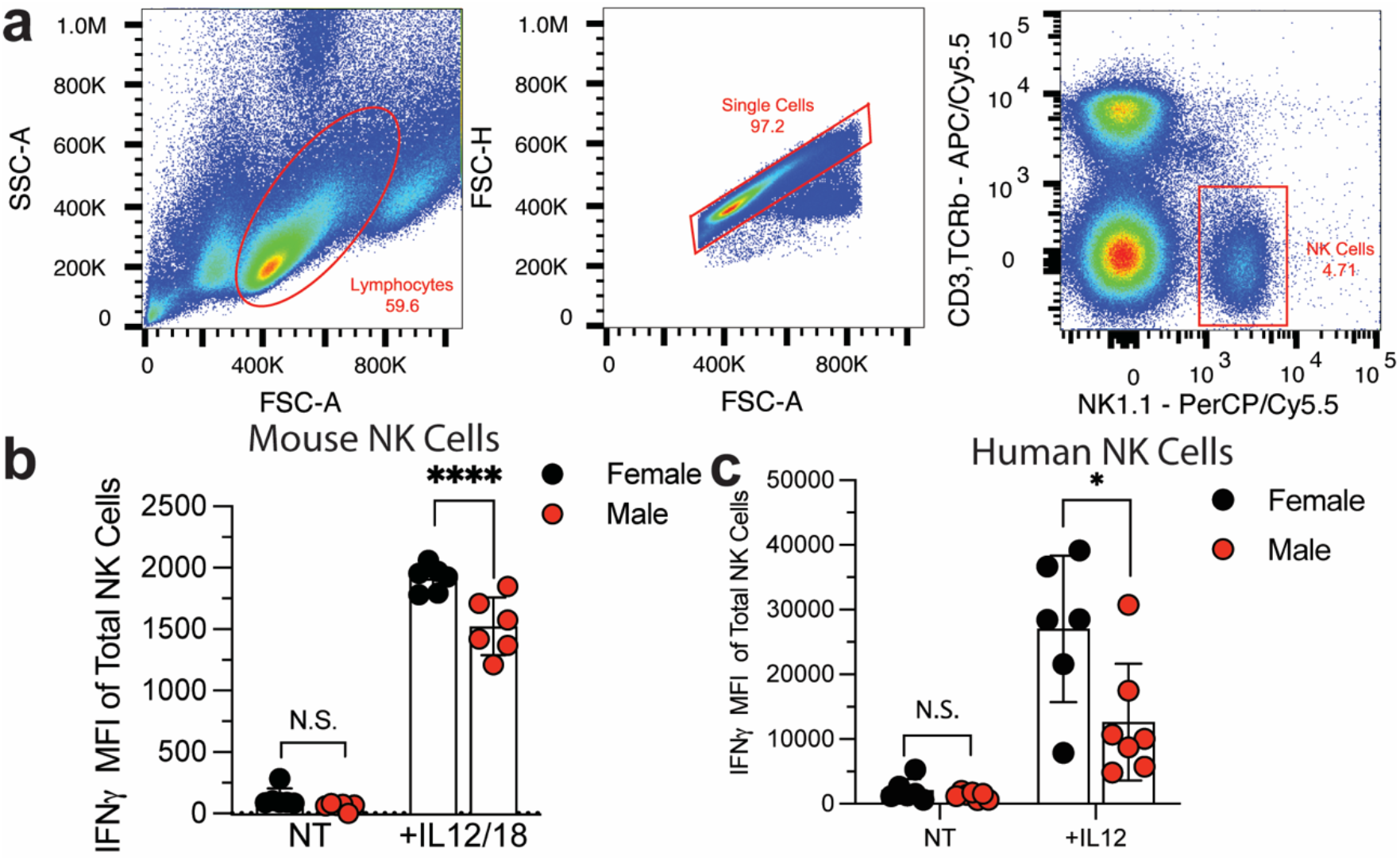
Male NK cells from mice and humans produce less IFN-γ than females in response to proinflammatory cytokines. **a)** Gating strategy for identification of NK cells (CD3^-^,TCR-β^-^ NK1.1^+^) in spleen of C57BL/6 mice. MFI of IFN-γ of NK cells from **b)** mouse stimulated with no treatment (NT) or IL-12 (20 ng/ml) and IL-18 (10 ng/ml) and **c)** human NK cells stimulated with no treatment (NT) IL-12 (10ng/ml). Samples were compared using paired two-tailed Student’s t test and data points are presented as individual samples with the mean ± SEM (N.S., Not Significant; *, p<0.05; ****, p<0.0001).

**Extended Data Figure 2:**
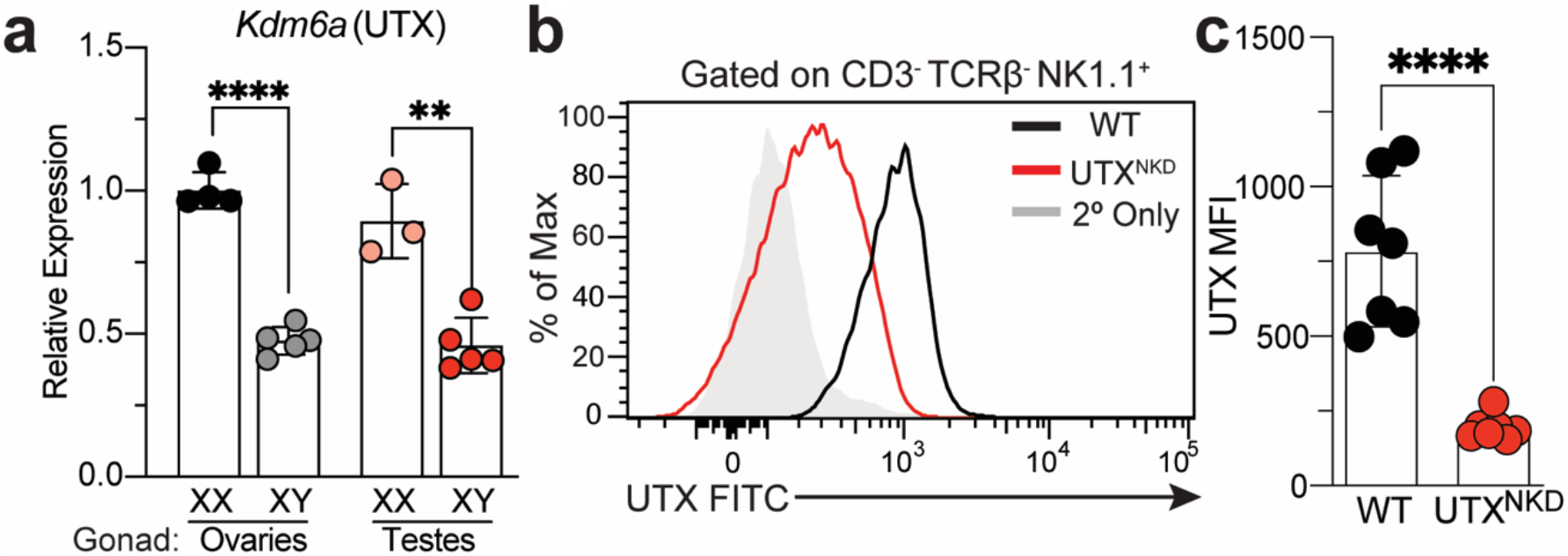
UTX expression in FCG and UTX^NKD^ mice. **a)** Relative expression of *Kdm6a* (UTX) by RT-qPCR from NK cell isolated from spleens of Four Core Genotype (FCG) mice (gonadal female XX n = 5, gonadal female XY n = 5, gonadal male XX n = 3, and gonadal male XY n = 5). **b)** Representative histogram plot and **c)** MFI of UTX protein expression in splenic NK cells (CD3^-^ TCRβ^-^ NK1.1^+^) in WT (*Kdm6a*^fl/fl^ x *Ncr1*^cre-^) and UTX^NKD^ (*Kdm6a*^fl/fl^ x *Ncr1*^cre+^) mice at steady state (n = 7). Samples were compared using paired two-tailed Student’s t test and data points are presented as individual mice with the mean ± SEM (**,p<0.01; ****, p<0.0001).

**Extended Data Figure 3:**
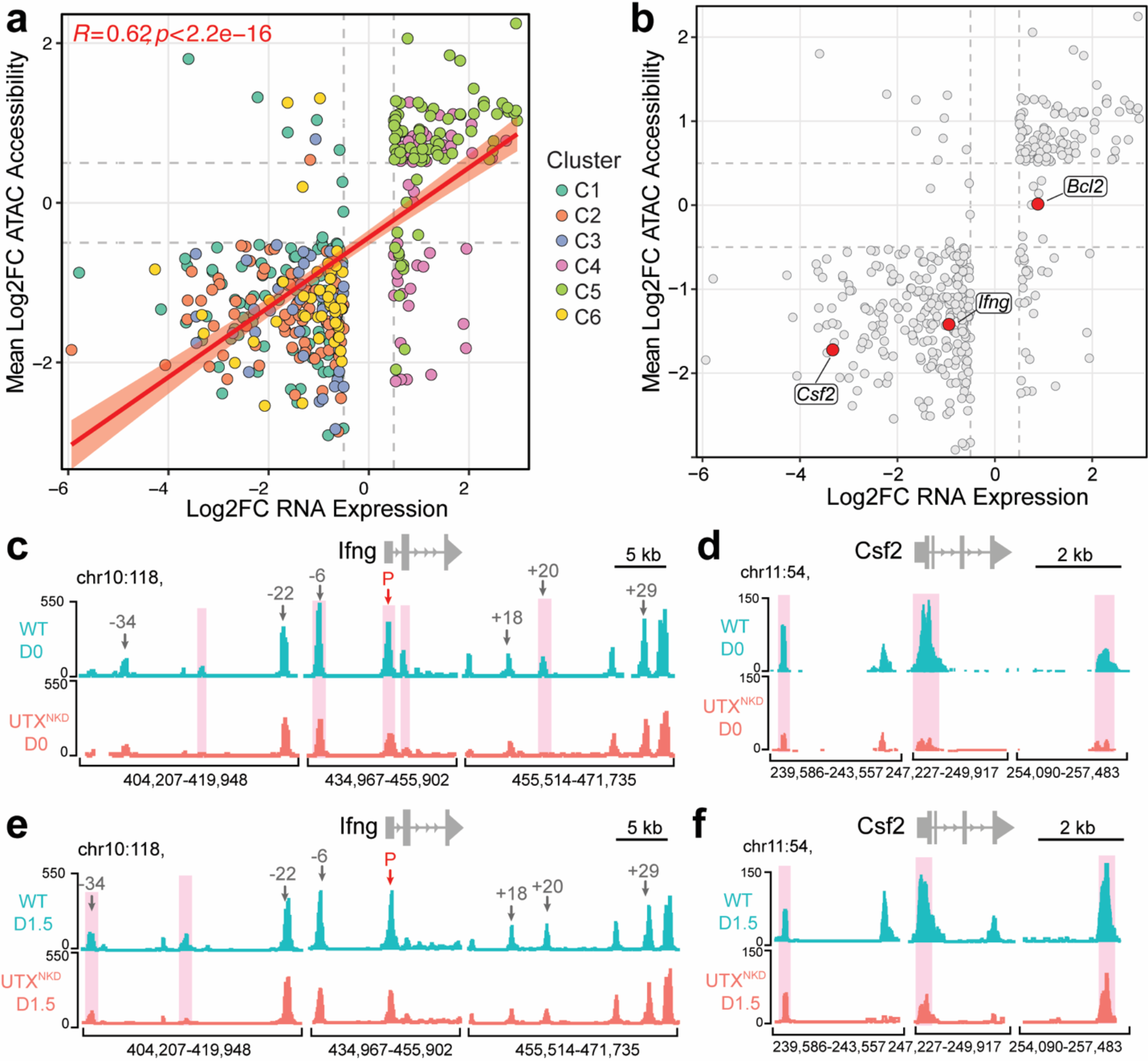
Integrative ATAC and RNA seq analysis reveal concomitant changes in chromatin accessibility and transcription mediated by UTX. a) Scatter plot highlighting genes that were differentially accessible and expressed (FDR and adjusted p-value < 0.05) colored by fuzzy c-means cluster (see Figure 4). Y-axis depicts mean log2 fold change of ATAC accessibility peaks and x-axis depicts log2 fold change of RNA-seq transcript levels. Best fit regression line (red) with standard error (light red ribbon). Positive correlation calculated by Spearman correlation of dataset (R=0.62, p<2.2×10^-16^). b) Specific genes of interest highlighted in red (*Ifng, Csf2, Casp3, Bcl2*) with known roles in NK cell gene programs. c-f) Representative tracks of ATAC-seq signal of genes involved in NK effector function c) *Ifng* and d) *Csf2* in NK cells isolated at rest (D0) and e) *Ifng* and f) *Csf2* 1.5 days post MCMV infection from WT:UTX^NKD^ mBMCs. Red highlighted areas represent significant differentially accessible regions (FDR and p-value < 0.05).

**Extended Data Figure 4:**
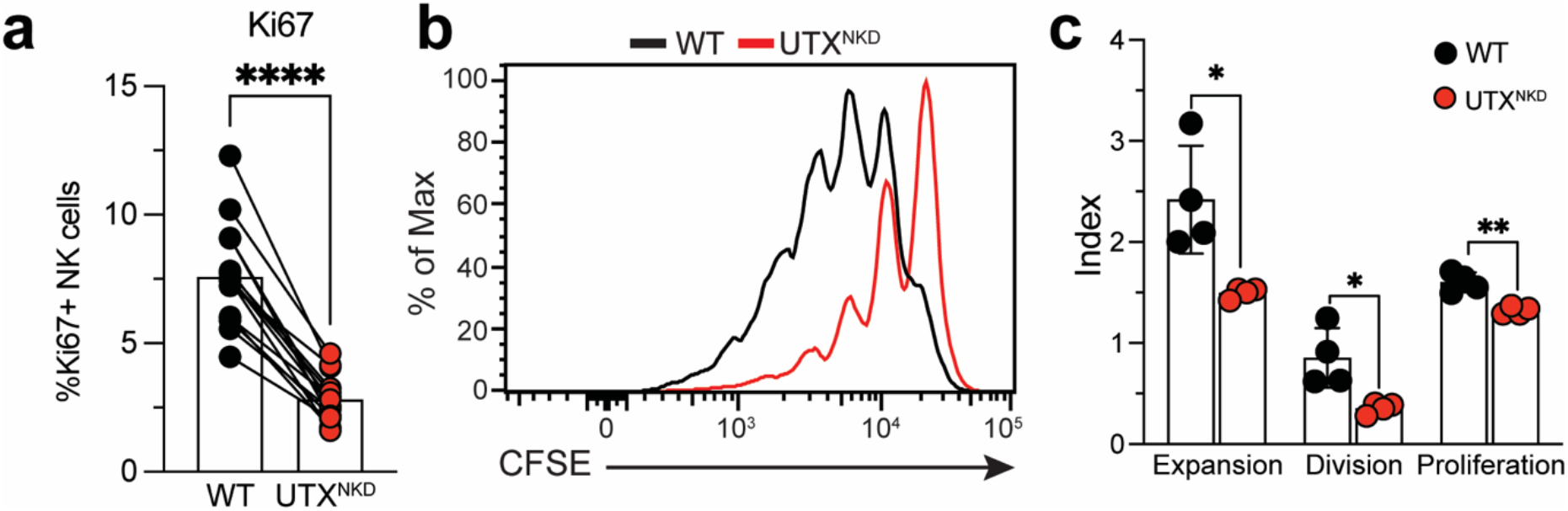
UTX deficient NK cells are less proliferative than WT. **a)** Percentage of Ki67^+^ total NK cells in blood of WT:UTX^NKD^ mBMCs (n=28). **b)** Representative flow cytometry plot and **c)** quantification of CFSE expansion, division and proliferation indexes of CFSE-labeled splenic NK cells from WT:UTX^NKD^ mBMCs stimulated *ex vivo* with IL-15 (50 ng/mL) for 4 days. Data represent 2-3 independent experiments. **a)** Samples were compared using paired and **b-c)** unpaired two-tailed Student’s t test and data points are presented as individual mice with the mean ± SEM (*, p <0.05; **, p <0.01).

**Extended Data Figure 5:**
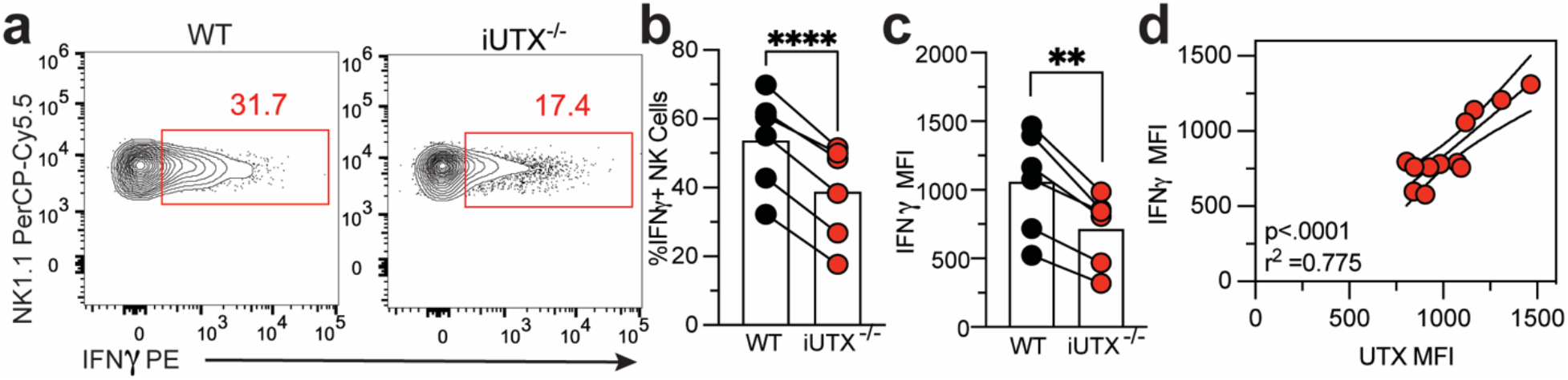
UTX regulates IFN-γ production in a dose-dependent manner. WT:UTX^-/-^ mBMCs were produced from WT (CD45.1^+^) and iUTX^-/-^ (CD45.2^+^) and treated with tamoxifen for 3 days (1mg/day) prior to MCMV infection. **g)** Representative flow cytometry plot and **h)** %IFN-γ^+^ and **i)** IFN-γ MFI of splenic NK cells at D1.5 post-MCMV infection of WT:iUTX^-/-^ mBMCs (n = 6). **j)** Correlation of IFN-γ vs. UTX MFI of total splenic NK cells (n = 12). Two-tailed correlation of XY data performed. Pearson’s r^2^ =0.775, and p <0.0001. Data are representative of 2-3 independent experiments. Samples were compared using paired two-tailed Student’s t test and data points are presented as individual mice with the mean ± SEM (**, p<0.01; ****, p<0.0001).

**Extended Data Figure 6:**
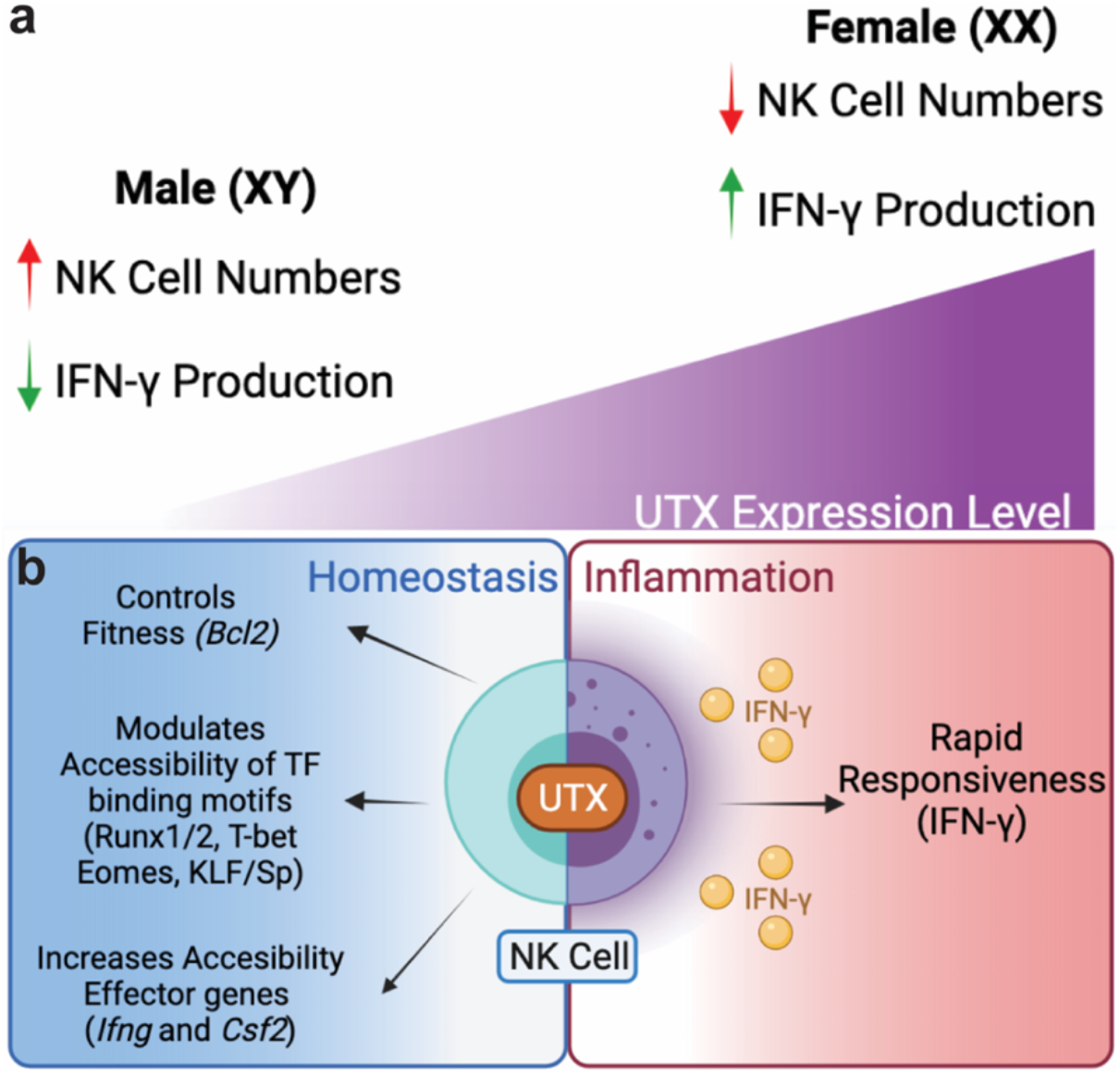
Schematic of UTX-mediated sex differences in regulation of NK cell homeostasis and effector function. **a)** Schematic of how differential UTX expression levels may underlie sexual dimorphism in NK cell composition and function. In male mice, lower UTX levels is associated with expansion of NK cell numbers and decreased NK cell IFN-γ production. **b)** Model in which UTX plays a role to control NK cell fitness and effector function. UTX controls 1) NK cell fitness through restricting Bcl-2 expression, 2) accessibility of TF binding motifs (Runx1/2, T-bet, Eomes, KLF/Sp), and 3) direct changes in chromatin accessibility of effector loci (*Ifng, Csf2*). Ultimately, UTX poises NK cells in an optimal epigenetic state to rapidly respond to viral infection by robust effector molecule production (IFN-γ) resulting in protection against viral infection.

## Contact for Reagent and Resource Sharing

Further information and requests for resources and reagents should be directed and will be fulfilled by the Corresponding Authors, Timothy O’Sullivan and Maureen Su

## Method Details

### Mice

Mice were bred at UCLA in accordance with the guidelines of the institutional Animal Care and Use Committee (IACUC). The following mouse strains were used in this study: C57BL/6 (CD45.2) (Jackson Labs, #000664), B6.SJL (CD45.1) (Jackson Labs, #002114), *Rosa26*^ERT2Cre^, *Ncr1*^Cre^, *Kdm6a*^fl/fl^ and FCG mice. For experiments with gonadectomy, procedure was performed by Jackson Laboratories Surgical Services. For experiments in UTX^NKD^ mice, only female mice were used to control for Y chromosome and sex hormone independent effects. Thus, experiments were conducted using 6-8 week old age-matched females in accordance with approved institutional protocols. For comparisons between male and female WT, we used 6-8 weeks age-matched littermates. Mixed bone marrow chimeras (mBMCs) were generated by depleting host CD45.1 x CD45.2 mice by intraperitoneal (i.p.) injection of busulfan (1mg/mL) at 20mg/kg for 3 consecutive days, which were then reconstituted 24 hours later with various mixtures of bone marrow cells from WT (CD45.1) and knockout (CD45.2) donor mice in the presence of an anti-NK1.1 antibody (at 1 mg/ml; clone: PK136) to deplete any remaining mature NK cells.

#### MCMV infection

MCMV (Smith) was serially passaged through BALB/c hosts three times, and then salivary gland viral stocks were prepared with a dounce homogenizer for dissociating the salivary glands of infected mice 3 weeks after infection. Experimental mice in studies were infected with MCMV by i.p. injection of 7.5 x 10^3^ plaque-forming units (PFU) in 0.5 mL of PBS. Mice were monitored and weighed daily and sacrificed when body weight dropped over 20% from initial weight.

#### Isolation and enrichment of mouse NK cells

Mouse spleens, livers, lungs, and blood were harvested and prepared into single cell suspensions as described previously^57^. Splenic single cell suspensions were lysed in red blood cell lysis buffer and resuspended in EasySep™ buffer (Stemcell). To avoid depleting Ly6C^+^ NK cells we developed a custom antibody cocktail as follows: splenocytes were labeled with 10 μg per spleen of biotin conjugated antibodies against CD3 (17A2), CD19 (6D5), CD8 (53-6.7), CD88 (20/70), Ly6G (1A8), SiglecF (S17007L), TCRβ (H57-597), CD20 (SA275A11), CD172a (P84) and magnetically depleted from total splenocyte suspensions with the use of anti-biotin coupled magnetic beads (Biolegend)^58^.

#### Ex vivo stimulation of lymphocytes

∼5 x 10^5^ NK cells were stimulated for 5 hours in complete RPMI media (RPMI 1640 + 25 mM HEPES + 10% FBS, 1% L-glutamine, 1% 200 mM sodium pyruvate, 1% MEM-NEAA, 1% penicillin-streptomycin, 0.5% sodium bicarbonate, 0.01% 55 mM 2-mercaptoethanol), Brefeldin A (1:1000; BioLegend) and Monensin (2uM; BioLegend) with or without recombinant mouse IL-12 (20 ng/ml; Peprotech) and recombinant mouse IL-18 (10ng/ml; Peprotech). Cells were cultured in complete RPMI media alone as a negative control (No Treatment or NT).

#### Human NK cell culture and Stimulation

Human peripheral blood mononuclear cells (PBMCs) from anonymous healthy donors were obtained from leukoreduction filters after platelet apheresis from the UCLA Virology Core. NK cells were isolated using the EasySep Human NK Cell Isolation Kit (Stem Cell Technologies) following manufacturer instructions. Following isolation, cells were maintained in 24-well G-Rex plates (Wilson Wolf) in NK MACS media (Miltenyi Biotech) supplemented with human IL-2 (100 IU/mL, Peprotech) and human IL-15 (20 ng/mL, Peprotech) at a plating density of 5×10^6^ cells per well. For cytokine stimulation, 14 d IL-2/IL-15 activated human NK cells were plated with K562 leukemia cells at an effector:target (E:T) ratio of 2.5:1 in addition to human IL-2 (100 IU/mL, Peprotech), human IL-15 (20 ng/mL, Peprotech), human IL-12 (10 ng/mL, Peprotech), and/or human IL-18 (100 ng/mL, Peprotech) in complete RPMI media (Thermo Fisher). NK cells were stimulated with cytokines for 16 h before analysis by flow cytometry.

#### Proliferation assays

CellTrace™ CFSE (Thermo) stock solution was prepared per the manufacturers’ instructions and diluted at 1:10,000 in 37C PBS. Isolated NK cells were incubated in 0.5mL of diluted CFSE solution for 5 minutes at 37C. The solution was quenched with 10X the volume of complete RPMI media. Cells were then washed and plated with 50 ng/ml of recombinant mouse IL-15 (Peprotech) and cultured for 4 days to assess proliferation. Flow cytometry was used to quantify CFSE dilution 4 days post-stimulation. Samples were compared to levels of CFSE labeled on Day 0. FlowJo’s proliferation tool was used to model and measure number of divisions in addition to expansion, division, and proliferation indices.

#### Flow Cytometry and Cell Sorting

Cells were analyzed for cell-surface markers using fluorophore-conjugated antibodies (BioLegend, eBioscience). Cell surface staining was performed in FACS Buffer (2% FBS and 2 mM EDTA in PBS) and intracellular staining was performed by fixing and permeabilizing using the eBioscience Foxp3/Transcription Factor kit for intranuclear proteins or BD Cytofix/Cytoperm kits for cytokines. Flow cytometry was performed using the Attune NxT Acoustic Focusing cytometer and data were analyzed with FlowJo software (TreeStar). NK cells were identified as CD3^-^ TCRβ^-^ NK1.1^+^ cells: see gating strategy in Extended Data Fig. 1. Cell surface and intracellular staining was performed using the following fluorophore-conjugated antibodies: CD45.1 (A20), CD45.2 (104), NK1.1 (PK136), TCRβ (H57-597), CD3 (17A2), IFN-γ (XMG1.2), Ly6C (HK1.4), BCL2 (BCL/10C4), UTX (N2C1 - GeneTex), Goat anti-rabbit H&L (Abcam - ab6717), BIM (c34c5), CD90 (30-H12), Ki-67 (16A8). Isolated splenic NK cells were sorted using Aria-H Cytometer to > 95% purity.

#### RNA-seq and ATAC-seq library construction and analysis

RNA was isolated from the cells using RNeasy Mini kit (Qiagen) and used to generate RNA-seq libraries followed by sequencing using Illumina HighSeq 4000 platform (single end, 50bp). The reads were mapped with HISAT2 (version 2.2.1) to the mouse genome (mm10). The counts for each gene were obtained by HtSeq^59^, in print, online at doi:10.1093/bioinformatics/btu638). Differential expression analyses were carried out using DESeq2^60^ (version 1.24.0) with default parameters. Genes with adjusted p value <0.05 were considered significantly differentially expressed. Sequencing depth normalized counts were used to plot the expression values for individual genes.

ATAC-seq libraries were produced by the Applied Genomics, Computation, and Translational Core Facility at Cedars Sinai in the following manner: 50,000 cells per sample were lysed to collect nuclei and treated with Tn5 transposase (Illumina) for 30 minutes at 37°C with gentle agitation. The DNA was isolated with DNA Clean & Concentrator Kit (Zymo) and PCR amplified and barcoded with NEBNext High-Fidelity PCR Mix (New England Biolabs) and unique dual indexes (Illumina). The ATAC-Seq library amplification was confirmed by real-time PCR, and additional barcoding PCR cycles were added as necessary while avoiding overamplification. Amplified ATAC-Seq libraries were purified with DNA Clean & Concentrator Kit (Zymo). The purified libraries were quantified with Kapa Library Quant Kit (KAPA Biosystems) and quality assessed on 4200 TapeStation System (Agilent). The libraries were pooled based on molar concentrations and sequenced on an Illumina HighSeq 4000 platform (paired end, 100bp).

ATAC-seq fastq files were trimmed to remove low-quality reads and adapters using Cutadapt^61^ (version 2.3). The reads were aligned to the reference mouse genome (mm10) with bowtie2^62^ (version 2.2.9). Peak calling was performed with MACS2^63^ (version 2.1.1). The peaks from all samples were merged into a single bed file, peaks from different samples that were closer than 10bp were merged into a single peak. HTseq^59^ (version 0.9.1) was used to count the number of reads that overlap each peak per sample. The peak counts were analyzed with DESeq2^60^ (version 1.24.0) to identify differentially accessible genomic regions. Peaks with adjusted p-value < 0.05 were considered significantly differentially accessible. The peak counts were visualized with Integrated Genome Browser, (version 9.1.8).

Fuzzy c-means clustering was used for both ATAC-seq and RNA-seq using significant (p-value and FDR <0.05, Log2FC +/- 0.5) normalized counts generated from DESeq2. MFuzz package (version 3.14) within R was used to perform this analysis into 6 clusters with a membership score of >0.5. The differentially accessible ATAC peaks were analyzed using the findMotifsGenome.pl function from HOMER^37^ (version 4.9.1) of each cluster to identify enriched cis-regulatory motifs of transcription factors. Pathway analysis of clustered RNA-seq data was performed using g:Profiler using the g:GOSt function. Top relevant pathways were selected from KEGG Biological Pathways and Gene Ontology Pathways (Biological Processes and Molecular Function).

#### Statistical Analyses

For graphs, data are shown as mean ± SEM, and unless otherwise indicated, statistical differences were evaluated using a Student’s t-test. For Kaplan-Meir survival curve, samples were compared using the Log-rank (Mantel-Cox) test with correction for testing multiple hypotheses. A p-value < 0.05 was considered significant. Graphs were produced and statistical analyses were performed using GraphPad Prism and ggplot2 library in R. Spearman Correlation on best fit regression line was performed using ggpubr library in R.

#### Data Availability

Sequencing datasets are accessible from GEO with accession number GSE185065. Data can be accessed by reviewers using the access token ilqlqswavvqhtir.

